# Stable isotope fingerprinting can directly link intestinal microorganisms with their carbon source and captures diet-induced substrate switching *in vivo*

**DOI:** 10.1101/2024.12.10.627769

**Authors:** Angie Mordant, J. Alfredo Blakeley-Ruiz, Manuel Kleiner

## Abstract

Diet has strong impacts on the composition and function of the gut microbiota with implications for host health. Therefore, it is critical to identify the dietary components that support growth of specific microorganisms *in vivo*. We used protein-based stable isotope fingerprinting (Protein-SIF) to link microbial species in gut microbiota to their carbon sources by measuring each microbe’s natural ^13^C content (δ^13^C) and matching it to the ^13^C content of available substrates. We fed gnotobiotic mice, inoculated with a 13 member microbiota, diets in which the ^13^C content of all components was known. We varied the source of protein, fiber or fat to observe ^13^C signature changes in microbial consumers of these substrates. We observed significant changes in the δ^13^C values and abundances of specific microbiota species, as well as host proteins, in response to changes in ^13^C signature or type of protein, fiber, and fat sources.

Using this approach we were able to show that upon switching dietary source of protein, fiber, or fat (1) some microbial species continued to obtain their carbon from the same dietary component (e.g., protein); (2) some species switched their main substrate type (e.g., from protein to carbohydrates); and (3) some species might derive their carbon through foraging on host compounds. Our results demonstrate that Protein-SIF can be used to identify the dietary-derived substrates assimilated into proteins by microbes in the intestinal tract; this approach holds promise for the analysis of microbiome substrate usage in humans without the need of substrate labeling.

**Significance:** The gut microbiota plays a critical role in the health of animals including humans, influencing metabolism, the immune system, and even behavior. Diet is one of the most significant factors in determining the function and composition of the gut microbiota, but our understanding of how specific dietary components directly impact individual microbes remains limited. We present the application of an approach that measures the carbon isotope “fingerprint” of proteins in biological samples. This fingerprint is similar to the fingerprint of the substrate used to make the proteins. We describe how we used this approach in mice to determine which dietary components specific intestinal microbes use as carbon sources to make their proteins. This approach can directly identify components of an animal’s diet that are consumed by gut microbes.

## Introduction

Interactions between the intestinal microbiota and diet play key roles in health and disease (1). For example, short-chain fatty acids derived from dietary fiber and protein fermentation by the gut microbiota play a role in mucus layer formation (2, 3) and are anti-inflammatory (4), while fermentation of specific amino acids derived from proteins also produce toxins like putrescine, ammonia, and hydrogen sulfide which are detrimental to host health (5, 6). Oftentimes the key assumption made in studies investigating the effects of diet on the gut microbiota is that the microbes that respond to a diet consume specific dietary components and that this drives changes in their abundance (7–9). Correlations between diet changes and taxon abundances, however, could also be due to other causes, such the effect of antimicrobial factors in foods (10), cross-feeding on byproducts from other species that consume dietary components (11), or switching between use of dietary substrates and host foraging (12, 13). Since microbiota response to diet can have major health consequences, tools to determine the use of particular dietary components by specific microbiota members are urgently needed.

Several approaches can infer specific diet-derived substrates consumed by intestinal microbes. These approaches have used individual microbes or defined communities in gnotobiotic animals in combination with gene expression analyses and *in vitro* growth of bacteria on dietary components (12, 14); strains in which genes for use of specific substrates were knocked out (15) or knocked in (16); or stable isotope probing (SIP) using labeled substrates and measuring the ratio of stable carbon isotope (i.e expressed as δ^13^C) in cellular components (17). The δ^13^C of cellular components is a particularly powerful method because it is the only method that can directly identify the nutrient sources of different organisms by comparing the isotopic ratio of an organism’s cellular components (e.g., protein, DNA, lipids) with the isotopic ratio(s) of the nutrient source(s) the organism used to build those components (18–22). Already, insights into nutrient flows in gut microbiota have been obtained by SIP approaches, where labeled compounds are injected into the bloodstream or provided as dietary components to the host and then incorporation of the labeled compound is determined by observing changes in the isotopic signatures of DNA, proteins, or metabolites from the host or microbiota. For example, ^15^N- and ^13^C-labeled threonine injected into the bloodstream of mice revealed variable incorporation of these labels, presumably from proteins secreted into the gut, by microbial taxa in the intestine using a high-resolution secondary ion mass spectrometry (NanoSIMS) approach, which revealed variable preferences for host versus dietary proteins across different microbial taxa (23, 24). In addition, oral or injected provision of a wide array of different labeled proteins and metabolites to healthy mice fed a standard chow diet revealed the nutrient sources for different microbes and their associated metabolites (17). For example, short-chain fatty acids were shown to primarily be produced by the fermentation of fiber as opposed to amino acids, and different bacterial taxa were shown to favor dietary protein, urea, or host protein as their source of nitrogen, but specifically did not use circulating amino acids. While these SIP studies provide tremendous insight into the steady state preferences of bacterial taxa for specific substrates, they are limited by the need to deliver labeled compounds to the intestine, which limits SIP’s future useability in humans, but also makes it difficult to explore the nuance of preference for different sources of macronutrients (e.g., different sources of fiber, protein, and fat) by bacteria in the intestine.

Mass spectrometry based shotgun metaproteomics on its own can provide insight into bacterial preferences for different macronutrients because it is able to capture how gene expression in specific microbes responds to changes in host diet. For example, in gnotobiotic mice it was shown that switching fiber source changes the expression of genomic regions encoding glycoside hydrolases in *Bacteroides* species (25) and more recently we showed that *Bacteroides thetaiotaomicron* gene expression changes in the presence of different dietary protein sources (26). This approach tells us how the nutrient sources are affecting the gene expression of the specific organisms and gives us some hints as to what nutrients the microbes are consuming, but fails to provide direct evidence for which nutrient sources the bacteria are actually incorporating.

We recently developed a metaproteomics method (protein-SIF) for measuring the **<u>natural</u>**stable carbon isotope fingerprints (δ^13^C) of specific species in a microbial community. Living organisms carry their own distinct carbon isotopic signature based on their carbon source (substrate/food) and this signature, or stable isotope fingerprint (SIF), can be used to infer the organism’s carbon source(s), as has been done in field ecology studies (18). Our protein-SIF approach for microbial communities was validated and tested in a case study using a gutless worm, which revealed new insights into the carbon sources of the worm’s microbial symbionts (19). Herein we describe our application of protein-SIF to gnotobiotic mice, colonized with a defined bacterial community, that were fed a series of controlled diets that varied in the type of available protein, fiber, or fat. Our results suggest that protein-SIF provides direct evidence for substrate preferences of specific members of a microbial community without the need for labeled compounds.

## Materials and Methods

### Overall experimental design

We conducted two experiments to investigate the assimilation of dietary macronutrients by intestinal bacteria in gnotobiotic mice colonized with a 13 member defined community (12)(**Table 1**; **Figure 1A**). In Experiment 1 (n=5) we fed gnotobiotic mice defined diets differing only in the source of dietary protein (egg white protein, casein, or soy protein). In Experiment 2 (n=6) we fed gnotobiotic mice defined diets that differed either only in their source of dietary fiber (cellulose, inulin, or corn fiber) or in their source of dietary fat (corn oil, soybean oil, or sunflower oil). All of these nutrients had distinct natural isotopic signatures (**Figure 1B**). Each diet was fed for one week and a fecal sample was taken from each mouse at the end of the week. Fecal samples were stored in nucleic acid preservation (NAP) buffer upon collection and then frozen at -80°C within hours of collection (27). We also collected baseline samples prior to transitioning to the defined diets when the mice had been eating a standard chow diet (Lab diet 5010) for 21 days after colonization and again 1 week after returning to the standard chow at the end of each experiment. We measured all the samples using a protein-SIF metaproteomic approach (**Figure 1C**) (19).

**Figure 1:**
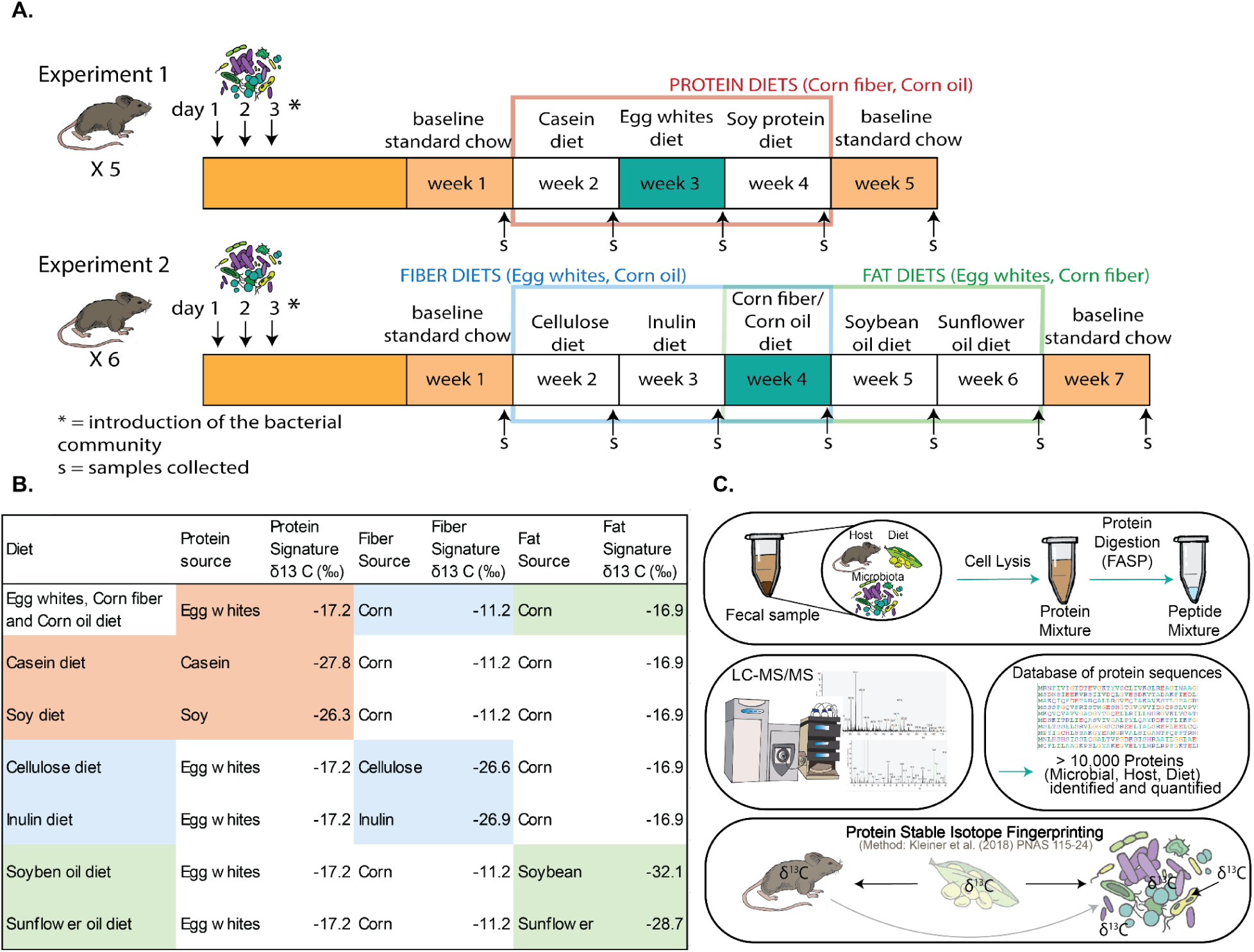
Overview of experimental design and procedure. A. Timeline and design of the experiment. The diet in the blue weeks (week 3 in Exp1 and week 4 in Exp2) was the same exact diet. B. Natural carbon isotopic signatures (δ^13^C values) of the macronutrients used in the diets. Signatures of dietary components were averaged between two replicates measured by EA-IRMS (measurement uncertainties of ± 0.42 ‰ or less; ± 0.13 ‰ averaged uncertainty). C. Overview of the metaproteomics protein-SIF approach.

**Table 1.** Organisms/strains of the defined community.

| Species | Strain | Original Source (before laboratory of Dr. Eric Martens) | Phylum |
| --- | --- | --- | --- |
| *Bacteroides ovatus* | 1896, Type strain | DSMZ | Bacteroidota |
| *Bacteroides uniformis* | 8492 | ATCC | Bacteroidota |
| *Bacteroides thetaiotaomicron* | 2079, Type strain | DSMZ | Bacteroidota |
| *Bacteroides caccae* | 19024, Type strain (ATCC43185) | DSMZ | Bacteroidota |
| *Barnesiella intestinihominis* | YIT11860 (JCM15079) | ATCC | Bacteroidota |
| *Roseburia intestinalis* | 14610, Type strain L1-82 | DSMZ | Bacillota |
| *Eubacterium rectale* | 17629, A1-86 | DSMZ | Bacillota |
| *Faecalibacterium prausnitzii* | 17677, A2-165 | DSMZ | Bacillota |
| *Marvinbryantia formatexigens* | 14469, Type strain I-52 | DSMZ | Bacillota |
| *Clostridium symbiosum* | 934, Type strain, designation 2 | DSMZ | Bacillota |
| *Collinsella aerofaciens* | 3979, Type strain | DSMZ | Actinomycetota |
| *Escherichia coli* | HS | ATCC | Pseudomonadota |
| *Akkermansia muciniphila* | 22959, Type strain, Muc | DSMZ | Verrucomicrobiota |

### Gnotobiotic models

We used 11 germ-free C57BL/J6 mice (5 in Experiment 1 and 6 in Experiment 2, all females). The NCSU Gnotobiotic core supplied and housed the mice. The mice were housed in groups of 3 throughout the experiment, except for one cage, which had 2 mice. All animal experiments followed protocols approved by the Institutional Animal Care and Use Committee (IACUC) of North Carolina State University.

The germ-free mice were colonized with the defined community (Table 1) at 9 - 11 weeks of age with freshly prepared bacterial inocula. Bacteria were grown in individual cultures in their respective media (Desai et al. 2016). The cultures were grown anaerobically in Hungate tubes.

The cultures were incubated at 37° C for 1-2 days depending on the strain until they reached optical densities (OD, absorbance at 600 nm) ranging from about 0.2 to >2. Bacterial cultures were mixed in equal volumes. We gavaged each mouse with 200 µL of the bacterial mixture for three consecutive days.

### Design of diets with known isotopic signatures

We designed 7 diets for this study based on the AIN93G (28) diet with minor modifications (**Suppl. Table 1**). All 7 diets contained the same source of starches and simple sugars: corn starch, maltodextrin, and sucrose. We included 3 groups of diets: “protein diets’’, “fiber diets’’, and “fat diets”. In each experiment, we compared 3 sources of protein, fiber, or fat. Every diet had one ingredient with a distinct isotopic signature from the other components in the diet. To design the diets, we measured the natural isotopic signatures of purified ingredients obtained from Envigo Teklad® and Amazon. We weighed the ingredients into tin capsules (∼ 1 mg per sample) and analyzed them by Elemental Analysis Isotope Ratio Mass Spectrometry (EA-IRMS) (**Figure 1B**). The ingredients were compared to a Vienna Pee Dee Belemnite (VPDB) standard using area under the curve calculations to determine the carbon isotope ratios.

We analyzed each purified ingredient in duplicate because IRMS measurements are highly robust (**Suppl. Table 2**). All defined diets were purchased from Envigo Teklad®. All defined diets were sterilized by gamma irradiation and vacuum packaged.

### Protein extraction and peptide preparation

Proteins for metaproteomics were extracted from samples of the seven diets (**Suppl. Table 1**), purified casein and purified egg white solids (to serve as a standard; see below), and feces from the eleven experimental mice at timepoints indicated in Figure 1A. We extracted the diets and purified ingredient samples in triplicate. Each replicate consisted of 100 mg of diet that we powdered down from diet pellets or 100 mg of a purified ingredient. For the fecal microbiome samples, we extracted one fecal pellet per mouse per diet. If applicable, we removed the NAP buffer from the samples by centrifugation at 21,000 x g for 5 min. We suspended the samples in 400 μl of SDT lysis buffer [4% (w/v) SDS, 100 mM Tris-HCl pH 7.6, 0.1 M DTT].

Cells were lysed by bead-beating in lysing matrix E tubes (MP Biomedicals) with a Bead Ruptor Elite (Omni International) for 5 cycles of 45 sec at 6.45 m/s with 1 min dwell time between cycles, followed by heating at 95° C for 10 min. Bead-beating was applied to the diet and the fecal samples but not the purified protein powders, which were heated at 95° C for 10 min. The lysates were centrifuged for 5 min at 21,000 x g to remove cell debris. We prepared peptides according to the filter-aided sample preparation (FASP) protocol (29). All centrifugations were performed at 14,000 x g. Samples were loaded twice onto 10 kDa MWCO 500 μl centrifugal filters (VWR International) by combining 60 μl of lysate with 400 μl of Urea solution (8 M urea in 0.1 M Tris/HCl pH 8.5) and centrifuging for 30 min. Filters were washed twice by applying 200 μl of urea solution followed by 40 min of centrifugation; 100 μl IAA solution (0.05 M iodoacetamide in Urea solution) was then added to filters for a 20 min incubation followed by centrifugation for 20 min. Filters were washed three more times by adding 100 μl of ABC (50 mM Ammonium Bicarbonate) followed by centrifugation for 20 min. Tryptic digestion was performed by adding 0.85 μg of MS grade trypsin (Thermo Scientific Pierce, Rockford, IL, USA) in 40 μl of ABC to the filters and incubating for 16 hours at 37° C. The tryptic peptides were eluted by adding 50 μl of 0.5 M NaCl and centrifuging for 20 min. Peptide concentrations were determined with the Pierce Micro BCA assay (Thermo Fisher Scientific) following the manufacturer’s instructions.

We mixed the peptides from the purified casein and the purified egg white solids in equal concentrations to create a casein and egg sample that was later used as an internal standard for the protein-SIF approach.

### LC-MS/MS

Samples were analyzed by 1D-LC-MS/MS as described in (27). The samples were blocked and randomized to control for batch effects. Alongside the fecal microbiome samples we included, at the beginning and end of the run sequence, two types of protein-SIF standards: the casein/soy protein standard mentioned above and peptides from human hair that were measured with EA-IRMS. Every sample was run as four consecutive technical replicates to increase the number of peptides available for protein-SIF. Only the human hair protein-SIF standard was run as a single replicate as it contained enough peptides. For each sample replicate, 600 ng of tryptic peptides were loaded with an UltiMate^TM^ 3000 RSLCnano Liquid Chromatograph (Thermo Fisher Scientific) in loading solvent A (2 % acetonitrile, 0.05 % trifluoroacetic acid) onto a 5 mm, 300 µm ID C18 Acclaim® PepMap100 pre-column and desalted (Thermo Fisher Scientific). Peptides were then separated on a 75 cm x 75 µm analytical EASY-Spray column packed with PepMap RSLC C18, 2 µm material (Thermo Fisher Scientific) heated to 60 °C via the integrated column heater at a flow rate of 300 nl min-1 using a 140 min gradient.

The analytical column was connected to a Q Exactive HF hybrid quadrupole-Orbitrap mass spectrometer (Thermo Fisher Scientific) via an Easy-Spray source. MS1 spectra were acquired by performing a full MS scan at a resolution of 60,000 on a 380 to 1600 m/z window. MS2 spectra were acquired using a data-dependent approach by selecting for fragmentation the 15 most abundant ions from the precursor MS1 spectra. A normalized collision energy of 25 was applied in the HCD cell to generate the peptide fragments for MS2 spectra. Other settings of the data-dependent acquisition included: a maximum injection time of 100 ms, a dynamic exclusion of 25 sec, and exclusion of ions of +1 charge state from fragmentation. About 60,000 MS/MS spectra were acquired per sample.

### Protein identification database

We constructed a protein sequence database for identifying proteins from the components of the fecal samples (i.e., the microbiota, the host, the dietary components, and potential contaminants) by downloading the relevant proteomes from UniProt (30, 31). We combined the genome of *Mus musculus* (UP000000589) with genomes of the strains used in this study (Table 1) and genomes to represent the origins of dietary proteins: *Gallus gallus* (Chicken UP000000539 - egg white protein)*, Glycine max* (Soybean UP000008827 - soy protein and soybean oil)*, Bos taurus* (Cow UP000009136 - casein)*, Zea mays* (Corn - corn fiber, corn oil, cornstarch)*, Helianthus annuus* (Sunflower - sunflower oil)*, and Beta vulgaris* (Sugar beet - sucrose). For the dietary and mouse proteomes, the protein sequences were clustered with an identity threshold of 95% using CD-HIT (Li and Godzik 2006). The protein sequences of the bacterial strains were not clustered. Also included in the database were sequences of common laboratory contaminants (http://www.thegpm.org/crap/). The database contains a total of 324,982 protein sequences and is available from the data submitted to the PRIDE repository.

### Protein identification and quantification

For peptide and protein identification, MS data were searched against the protein database using the Sequest HT node in Proteome Discoverer version 2.3.0.523 (Thermo Fisher Scientific) with the following parameters: digestion with trypsin (Full), maximum of 2 missed cleavages, 10 ppm precursor mass tolerance, 0.1 Da fragment mass tolerance and maximum 3 equal dynamic modifications per peptide. We considered the following dynamic modifications: oxidation on M (+15.995 Da), carbamidomethyl on C (+57.021 Da), and acetyl on the protein N terminus (+42.011 Da). Peptide false discovery rate (FDR) was calculated using the Percolator node in Proteome Discoverer, and only peptides identified at a FDR <5% were retained for protein identification. Proteins were inferred from peptide identifications using the Protein-FDR Validator node in Proteome Discoverer with a target FDR of 5%. We generated files of individual samples by combining the four replicate LC-MS/MS-produced files in the search. We used the resulting PSM file for the protein-SIF method.

### Protein stable isotope fingerprinting (protein-SIF)

We used the Calis-p 2.0 software (Kleiner et al., 2023) to determine stable isotopic fingerprints of the organisms in the samples. The Calis-p software requires two input files: raw spectral files produced by the LC-MS/MS, and the peptide-spectrum match (PSM) files containing the protein identifications and quantifications. Raw files were converted to mzML format using the MSConvertGUI tool via ProteoWizard (Chambers et al. 2012) with the following options: Binary encoding precision: 64-bit, Write index: checked, TPP compatibility: checked, Filter: Peak Picking, Algorithm: Vendor, MS Levels: 1. The PSM files were generated as described above, and input into the Calis-p software as tab-delimited text files.

Calis-p performs two main steps: isotopic pattern extraction and SIF computation. The software first filtered out “ambiguous” peptides of low identification confidence from the input PSM file. Then, for each remaining peptide identification, the software found the corresponding mass spectrum in the mzML and extracted the isotopic pattern. After some clean-up and filtering steps, the software compared the experimentally derived isotopic pattern to *in silico* derived isotopic patterns to infer δ^13^C values for the peptide. This step was repeated for all peptides.

Finally, the average δ^13^C of the peptides from an organism was used to estimate that organism’s signature. We filtered the results to retain only SIF values computed from at least 30 peptides, which is the threshold required by the Calis-p 2.0 software to accurately estimate an organism’s SIF.

We corrected for the offset introduced by the mass spectrometer using both the human hair standard and the casein/egg standard by collecting δ^13^C values for the standards obtained both by protein-SIF and EA-IRMS and calculating the offset between the two methods (19). We used the averaged offset value to correct the protein-SIF values of the organisms in the microbiome samples.

### Data analyses

We compared the SIF of each organism (**Table 1**) to the signatures of the dietary components fed to the mice. We looked for correspondence between signatures of organisms and signatures of dietary components to hypothesize which dietary constituents were assimilated by each organism. We also looked at how each organism’s SIF changed over time due to the different diet inputs to inform further data analyses. We identified significant differences (p < 0.05) using pairwise t-tests corrected for multiple hypotheses testing (Benjamini-Hochberg correction), computed in R version 4.0.2. We included the SIF values of the standard chow samples in the results to assess reproducibility. However, we did not know the signatures of the dietary constituents of the standard chow diet, and thus we did not test for significant differences between the standard chow and the defined diets. We prepared plots for organisms that had at least 3 data points from each of a minimum of two defined diets so that we could perform statistical analyses.

To estimate abundances of the different species, we applied the protein biomass method we developed previously (32). Briefly, we filtered the identified proteins to proteins that had at least 2 protein unique peptides (2PUP proteins), we then summed the peptide spectral matches (PSMs) of all of the 2PUP proteins for each organism, and then calculated a percent proteinaceous biomass for each organism. We tested for significant changes in relative abundances using a one-way ANOVA followed by a Tukey’s honestly significant difference (HSD) post hoc test, computed in R (version 4.0.2). We assigned letters to group means that are similar using the “rcompanion” package (version 2.4.1).

As a case study, to determine if the proteome of specific species could support the nutrient source hypotheses inferred from the protein-SIF data we calculated normalized protein abundance values (orgNSAF) for the proteins of *A. muciniphila*, *B. thetaiotaomicron*, and *M. formatexigens*. We then tested for significant differences in protein abundance between the different diets using an FDR-corrected ANOVA (q < 0.05). Plots were prepared in Origin 2018b, R pheatmap package, or in the Perseus software platform (version 1.6.12.0) (33), and compiled in Adobe Illustrator 2021.

## Results and Discussion

### Experiment 1: Changes in source of dietary protein impact isotopic signatures of intestinal microorganisms indicating their main carbon source

We fed gnotobiotic mice, colonized with a 13 member community (**Table 1**), a sequence of three defined diets that differed in their dietary protein source (casein, egg white, soy); our goal was to determine if these changes in dietary protein source affected the natural isotopic signature of intestinal microbes and the host, which would provide evidence for carbon source preferences.

The δ^13^C values of the dietary protein sources were -26.6‰ for casein, -17.2‰ for egg white, and -26.3‰ for soy (**Suppl. Table S2**). The isotopic signatures of all other dietary components were kept steady at -10.7‰ (cornstarch), -12.3‰ (sucrose), -11.2‰ (corn fiber), and -16.9‰ (corn oil). We switched the protein source every seven days after collecting fecal samples (**Figure 2A**). Our rationale was that a significant increase in the δ^13^C value of an organism between the casein and egg white diets followed by a decrease in δ^13^C value between the egg white and soy diets, i.e., a change in microbial δ^13^C value that tracked the diet’s change in δ^13^C value, would indicate that the organism uses dietary protein as a carbon source to make protein. We would not necessarily expect that the δ^13^C value of an organism becomes identical, or nearly identical, to that of a dietary component as an organism may use multiple substrates and carbon previously present in a cell will “dilute” the δ^13^C value of a newly used substrate. From each fecal sample we used metaproteomics to (1) extract microbial and host protein isotopic signatures using the protein-SIF approach (19); and (2) determine microbial community composition in terms of proteinaceous biomass (Kleiner et al., 2017).

**Figure 2:**
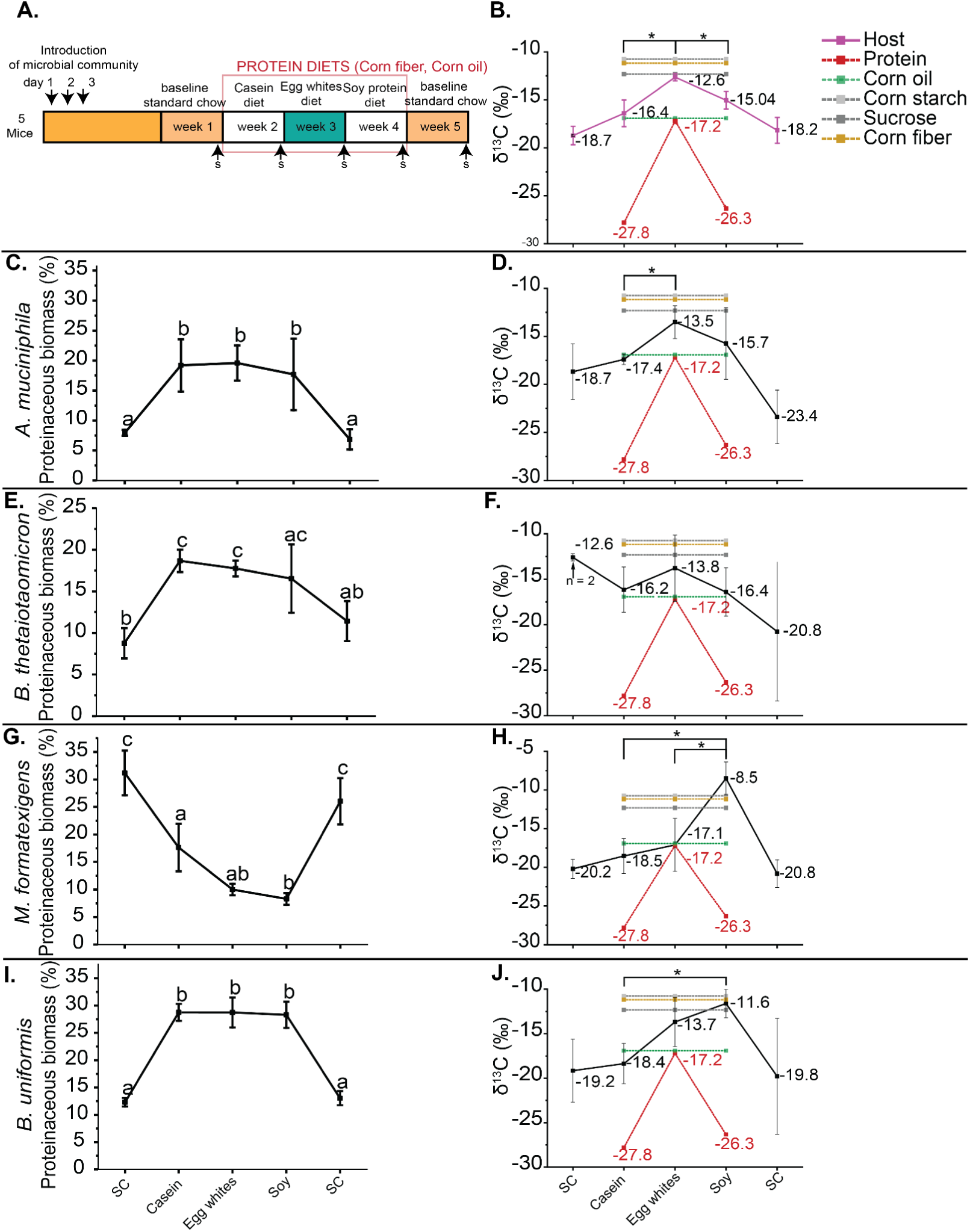
Changes in isotopic signatures of gut microbes in response to changes in the isotopic signature or source of dietary protein. (A) Overview of experiment 1 as described in Materials & Methods (Figure 1). (B, D, F, H, J) Protein-SIF δ^13^C values for mouse, *A. muciniphila*, *B. thetaiotaomicron*, *M. formatexigens* and *B. uniformis*. Displayed δ^13^C values for dietary protein sources, corn oil, corn starch, sucrose, and corn fiber were measured by isotope ratio mass spectrometry (IRMS). Significance is denoted by * and determined by T-tests corrected by BH (p < 0.05; n=5 unless otherwise stated). (C, E, G, I) Relative proteinaceous biomass of *A. muciniphila*, *B. thetaiotaomicron*, *M. formatexigens* and *B. uniformis* determined according to the method described by (32). Letters that do not overlap denote significantly different groups as determined by Tukey HSD (p < 0.05).

The protein-SIF signature of host proteins detected in the feces increased significantly in the egg white protein diet compared to the other two diets: the average mouse δ^13^C value on egg white was -12.6‰, compared to -16.4‰ and -15.0‰ on casein and soy protein, respectively, (BH corrected t-test p < 0.05)(**Figure 2B**). Because the change in δ^13^C value of the host proteins mirrors that of the diet, this indicates that dietary protein is a carbon source for the host.

We were able to determine δ^13^C values for four members of the defined community: *A. muciniphila*, *B. thetaiotaomicron*, *M. formatexigens* and *B. uniformis*. The abundance of *A. muciniphila* significantly (Tukey HSD p<0.05) increased when switching from the initial standard chow diet to the casein diet and did not change after the casein-to-egg white or egg white-to-soy switches (**Figure 2C**). The δ^13^C value of *A. muciniphila* significantly increased, to -13.5‰, in response to the egg white diet, indicating *A. muciniphila*’s metabolism responded to changes in dietary protein, mirroring the isotopic signature changes in the host (**Figure 2D**).

Since the host also responds to the dietary protein source in the same way and *A. muciniphila* is known to grow on host intestinal glycoproteins (mucins) (12, 34), this result is likely due to the use of host proteins as a carbon source by *A. muciniphila*. The abundance of *B. thetaiotaomicron* also increased after transitioning from standard chow to the casein diet and remained at a similar level with the egg white and soy diets (**Figure 2E)**. Although the δ^13^C value of *B. thetaiotaomicron* was higher under the egg white than the casein diet, this difference and other δ^13^C value comparisons were not statistically significant (**Figure 2F**). The δ^13^C values of *M. formatexigens* and *B. uniformis* were the highest under the soy protein diet, with some differences being statistically significant (**Figure 2H and 2J**). This was unexpected, since the δ^13^C value of soy protein is lower than the preceding egg white protein and therefore the expectation was that the δ^13^C values of these species would decrease if they were using protein as a carbon source or remain stable if not using protein as a carbon source. Instead, the increasing δ^13^C values suggest that *M. formatexigens* and *B. uniformis* transitioned from incorporating carbon from dietary or host protein to incorporating most carbon from the available carbohydrate sources (e.g. starch, sucrose, or corn fiber) when the diet was transitioned from casein to soy protein. Interestingly, the abundance of *M. formatexigens* decreased between the casein and soy protein diets (**Figure 2G**), while the abundance of *B. uniformis* did not change in response to dietary protein source (**Figure 2I**). Together these results indicate that the isotopic signatures of microbiota members and the host respond within a maximum of seven days after changing the diet. The results also indicate that changes in abundance are not necessarily driven by continued consumption of the same major substrate (i.e. protein), and that when the available substrate changes, e.g., from egg white protein to soy protein, the bacteria may preferentially consume fiber or fat. .

In addition to these bacteria for which we could get an isotopic signature, we were also able to measure the abundances of *B. ovatus, R. intestinalis, E. rectale, E. coli, B. caccae, B. intestinihominis, C. aerofaciens, and C. symbiosum* (**Suppl. Figure 1**)*. B. ovatus, B. intestinihominis,* and *E. rectale* significantly changed in abundance when transitioning between standard chow and the defined diets. *B. ovatus*, *E. coli*, *B. caccae*, *B. intestinihominis,* and *C. aerofaciens* significantly changed in abundance when transitioning between sources of dietary protein. Together these results show that dietary protein source alters the abundances of specific bacteria in the gut as we have previously shown in conventional mice (26).

### Experiment 2: Changes in source of fiber or fat alters the isotopic signatures of *B. thetaiotaomicron*, *M. formatexigens* and *A. muciniphila*

For experiment 2, we colonized germ-free mice with the same gnotobiotic community, but offered only egg white as the source of dietary protein. We first changed the fiber source each week, going from cellulose (-26.5‰), to inulin (-26.99‰), to corn fiber (-11.2‰) (**Figure 1A&B; Figure 3A**). We then kept the fiber isotopic signature constant by using corn fiber and changed the fat source each week from corn oil (-16.9‰), to soybean oil (-32.1‰), and to sunflower oil (28.7‰). Neither fiber nor fat source significantly altered the δ^13^C value of the mouse proteins, which hovered at approximately -11‰ (**Figure 3B**).

**Figure 3:**
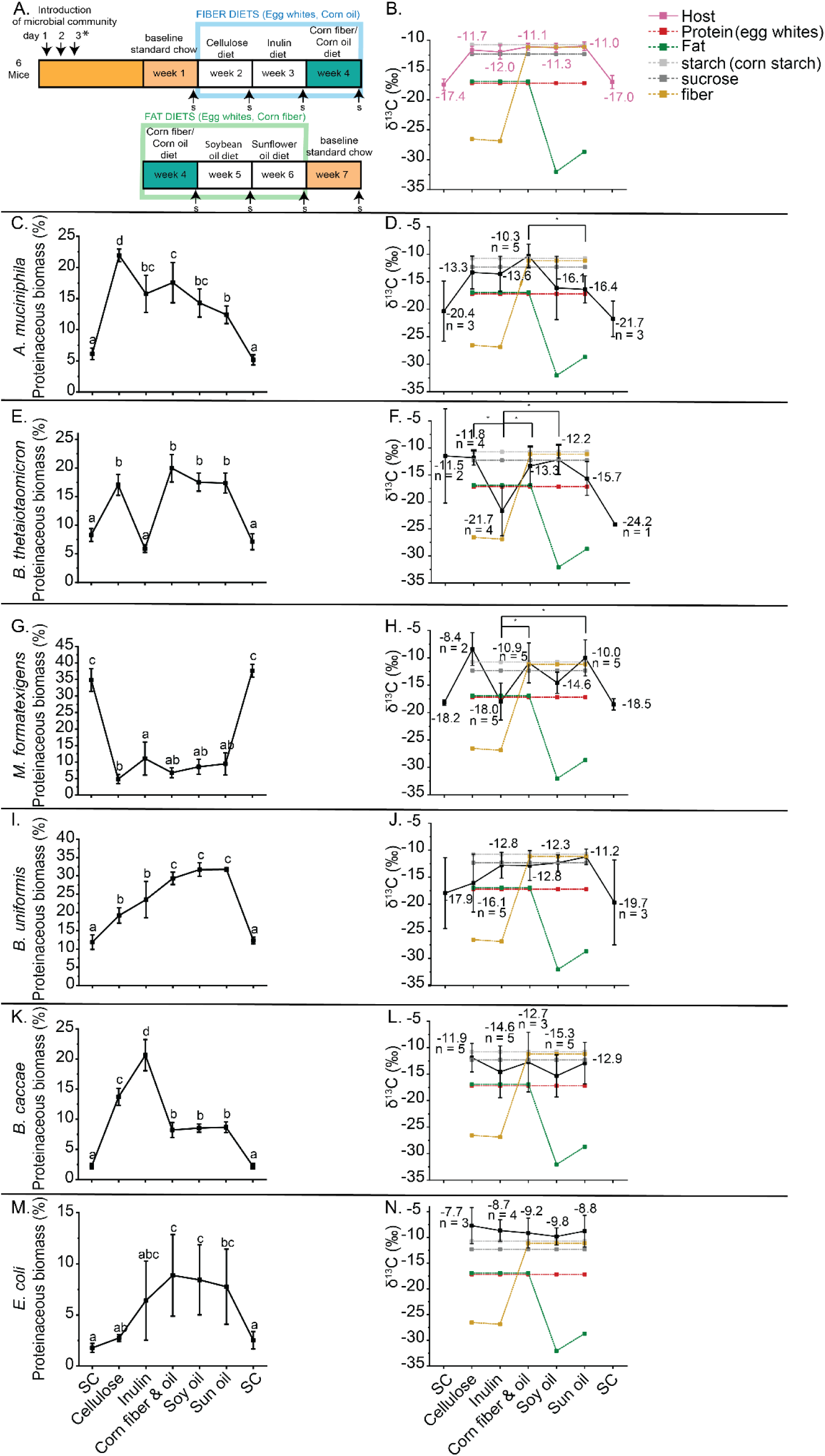
Changes in isotopic signatures of gut microbes in response to changes in the isotopic signature or source of dietary fiber or fat. A. Overview of Experiment 2 as described in Materials & Methods (Figure 1). (B, D, F, H, J, L, N) Protein-SIF δ^13^C values for mouse, *A. muciniphila*, *B. thetaiotaomicron*, *M. formatexigens*, *B. uniformis*, *B. caccae*, and *E. coli*, respectively. Displayed δ^13^C values for dietary fiber and fat sources, corn starch, sucrose, and egg white protein were measured by isotope ratio mass spectrometry (IRMS). Significance is denoted by * and determined by T-tests corrected by BH (p < 0.05; n=6 unless otherwise stated). (C, E, G, I, K, M) Relative proteinaceous biomass of *A. muciniphila, B. thetaiotaomicron, M. formatexigens and B. uniformis, B. caccae,* and *E. coli*. Letters that do not overlap denote significantly different groups as determined by Tukey HSD (p < 0.05).

The only microbe whose isotopic signature changed in response to fat source was *A. muciniphila*. Its δ^13^C value decreased steadily as it transitioned between the corn oil, soybean oil, and sunflower oil diets with a significant difference between the δ^13^C values under the corn oil diet and the sunflower oil diet (**Figure 3D**). Both soybean oil and sunflower oil have much lower δ^13^C values than corn oil. The abundance of *A. muciniphila* was also responsive to fiber and fat source (**Figure 3C**). Its abundance significantly increased from the standard chow to the defined diets, reaching its highest level under the cellulose diet. Additionally, the abundance of *A. muciniphila* significantly decreased in the sunflower oil diet relative to the cellulose, inulin, and corn fiber-corn oil diets.

Both *B. thetaiotaomicron* and *M. formatexigens* changed significantly in their abundances and isotopic signatures in response to changes in source of fiber. Interestingly, the abundance and isotopic signature of *B. thetaiotaomicron* significantly decreased when switching the diet from cellulose to inulin (**Figure 3E & F**). This was unexpected because cellulose and inulin had similar δ^13^C values and thus the change in *B. thetaiotaomicron* isotope signature cannot be explained by continued use of fiber as carbon substrate. The δ^13^C values of other potential substrates for *B. thetaiotaomicron* including starch, sucrose, and the host were much higher than the ones of cellulose and inulin. Taken together this suggests that the change in *B. thetaiotaomicron* isotopic signature was driven by a switch from one of these nutrients to inulin despite inulin not leading to an increase in the abundance of *B. thetaiotaomicron.* In the case of *M. formatexigens*, the bacterium had a lower δ^13^C value in the inulin diet relative to the corn fiber containing diets and trended towards a lower δ^13^C in the inulin diet relative to the cellulose diet (**Fig. 3F**). In this case, *M. formatexigens* significantly increased in abundance between cellulose and inulin suggesting that the transition towards incorporating inulin in this case increased the abundance of *M. formatexigens* (**Fig. 3E**).

Isotopic signatures for *B. uniformis*, *B. caccae*, and *E. coli* did not change in response to changes in fiber or fat sources (**Figure 3J**, 3L **and 3N**). All three of these bacteria, however, changed in abundance due to fiber source. *B. uniformis* (**Fig. 3I**) significantly increased in abundance between the cellulose and corn fiber-corn oil diets, *B. caccae* significantly increased in abundance between cellulose and inulin then significantly decreased in the presence of corn fiber (**Fig. 3 K**), and *E. coli* increased in abundance between the cellulose and corn fiber diets as well (**Fig. 3M**).

We were able to calculate the abundances of *B. ovatus*, *R. intestinalis*, *E. rectale*, *B. intestinihominis*, *C. symbiosum*, and *C. aerofaciens* (**Suppl. Figure 2**). None of these bacteria had a significant change in abundance due to fat source, but they all experienced significant changes in abundance due to fiber source suggesting that fiber source has a much greater impact on gut microbiota composition than fat source.

### Differential metaproteomic analysis reveals additional information about carbon source choices of specific species

We analyzed the metaproteomic data to identify gene expression changes of specific species in response to diet changes to gain additional insights into the nutrient/carbon sources of species with unexpected changes in their isotopic signatures. We focused on *M. formatexigens* in Experiment 1, *B. thetaiotaomicron* in the fiber component of Experiment 2, and *A. muciniphila* in the fat component of Experiment 2.

We identified 60 *M. formatexigens* proteins that significantly changed in abundance between casein, egg white, or soy protein (ANOVA, q < 0.05). Hierarchical clustering of samples using these proteins revealed 17 proteins that were more abundant in the soy diet relative to the other diets (**Figure 4**). Among these were two proteins that are components of a sugar transport ABC transporter, which suggests more sugar was being imported under the soy diet. Also increased in the soy diet was the abundance of *M. formatexigens’* glutamine synthetase, which is a protein that is upregulated in bacteria under nitrogen limitation, when it plays a major role in the incorporation of inorganic nitrogen (35). Together these results suggest that the soy diet-induced shift in *M. formatexigens’* isotopic signature was due to a switch from protein to sugar as the primary carbon source. It is unclear if the increase in glutamine synthetase expression was caused by generally lower nitrogen availability in the gut during the soy protein diet or by a need for *M. formatexigens* to acquire inorganic nitrogen for *de novo* amino acid synthesis when not using soy protein as a source of organic carbon and nitrogen.

**Figure 4:**
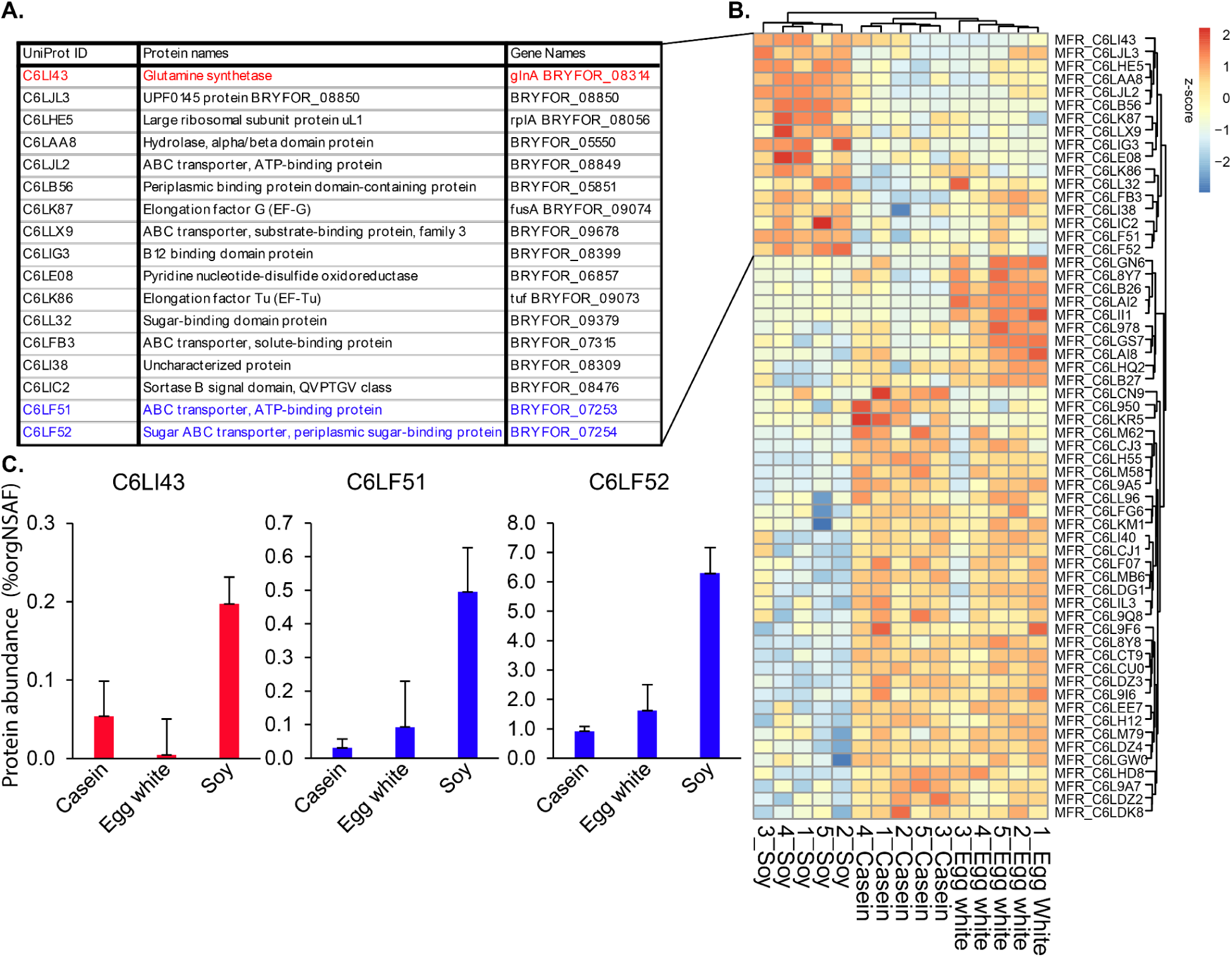
Hierarchical clustering of *M. formatexigens* proteins that significantly differed between casein, egg, white and soy protein. A. Table listing 17 proteins that were more abundant in the soy cluster. B. Hierarchical clustering of the z-score values of the 60 proteins that changed significantly in abundance between the three dietary protein sources (ANOVA, q < 0.05, n=5). C. Abundant proteins that potentially explain the increase in the δ^13^C value in the soy diet.(Figure 2H). Glutamine synthetase is highlighted in red; proteins from a Sugar ABC transporter gene neighborhood are highlighted in blue. Bars represent average% orgNSAF abundance and error bars represent the standard deviation.

We identified 147 *B. thetaiotaomicron* proteins whose abundances significantly differed between the cellulose, inulin, and corn fiber diets (ANOVA, q < 0.05). Using hierarchical clustering, these proteins separated into distinct inulin, cellulose, and corn fiber clusters. The inulin cluster was most distinct containing 41 proteins whose abundances increased in response to inulin relative to the other diets (**Figure 5**). Fifty of the 147 significantly different proteins, approximately ⅓, belong to polysaccharide utilization loci (PUL)(Grondin et al., 2017; Terrapon et al., 2018) (**Figure 6**). PULs are *Bacteroides* gene neighborhoods that encode all the enzymes needed to import and degrade a specific glycan structure. Six of these PUL-derived proteins, which were among the 41 proteins significantly increased under the inulin diet relative to the other diets, represent ⅔ of the 9 proteins encoded by the inulin- and levan-degrading PUL22 (All PUL numbers are from the lit-derived numbering in PULDB) (37, 38). The remaining PULs were elevated in cellulose or corn fiber relative to inulin. PUL66 is a starch degrading PUL, while PUL14, PUL19, PUL72, PUL73 PUL80, and PUL81 have been previously linked to degrading host protein glycosylations (14). We also recently linked PUL14, PUL72, and PUL80 to the degradation of egg white protein glycosylations, which have glycan structures similar to host intestinal mucin (26). Together with the observed changes in δ^13^C values under the inulin diet (**Figure 3F**), these results suggest that *B. thetaiotaomicron* transitioned from using starch and a combination of host and dietary proteins under the cellulose and corn fiber diets, to using inulin as its primary carbon source under the inulin diet.

**Figure 5:**
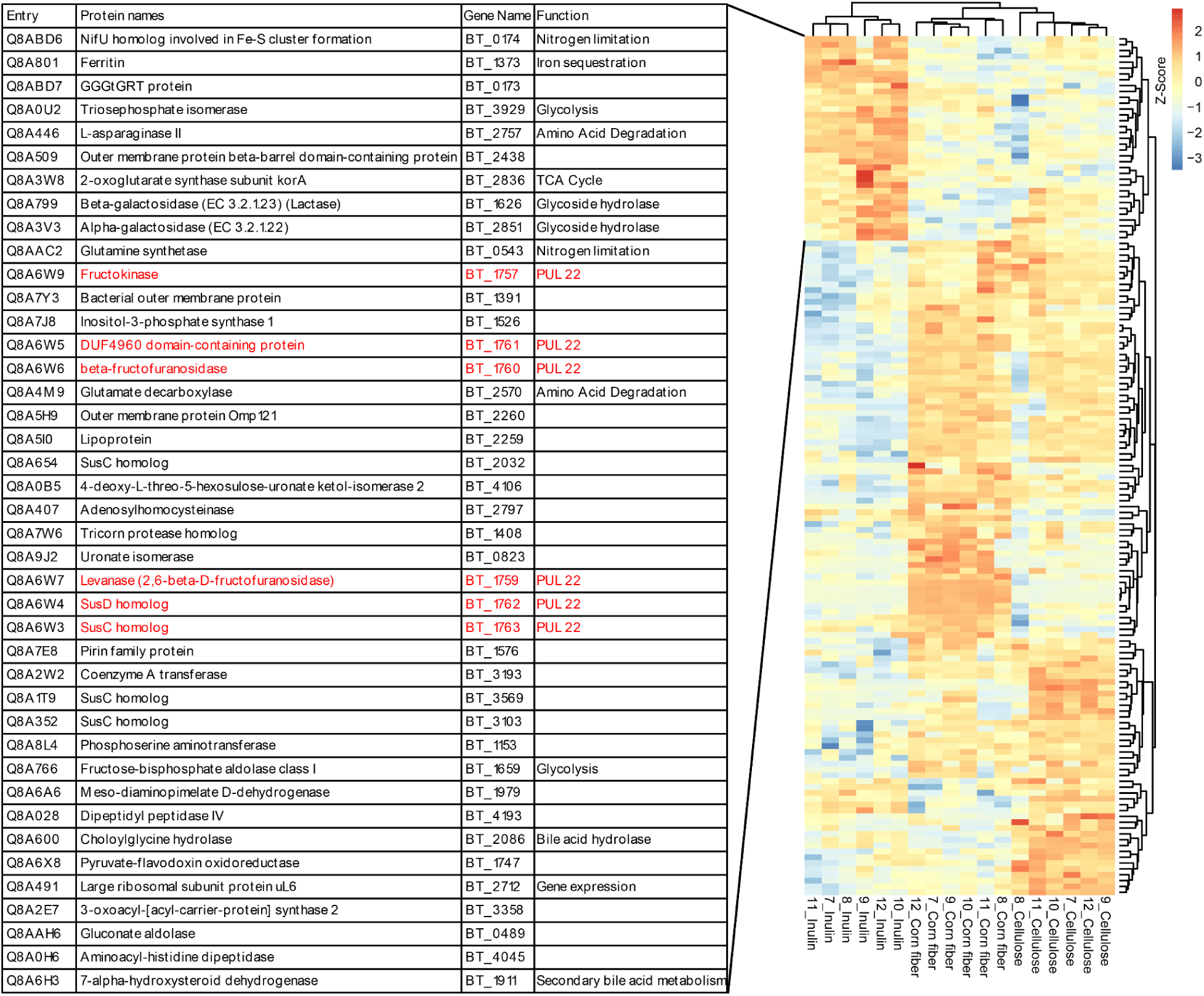
Hierarchical clustering of *B. thetaiotaomicron* proteins that significantly differed between cellulose, inulin, and corn fiber. Hierarchical clustering of the z-score values of the 147 proteins whose abundances changed significantly between the three dietary fiber sources (ANOVA, q < 0.05, n=5). The table represents the 41 proteins abundant in the inulin cluster. Proteins from PUL22 are highlighted in red.

**Figure 6:**
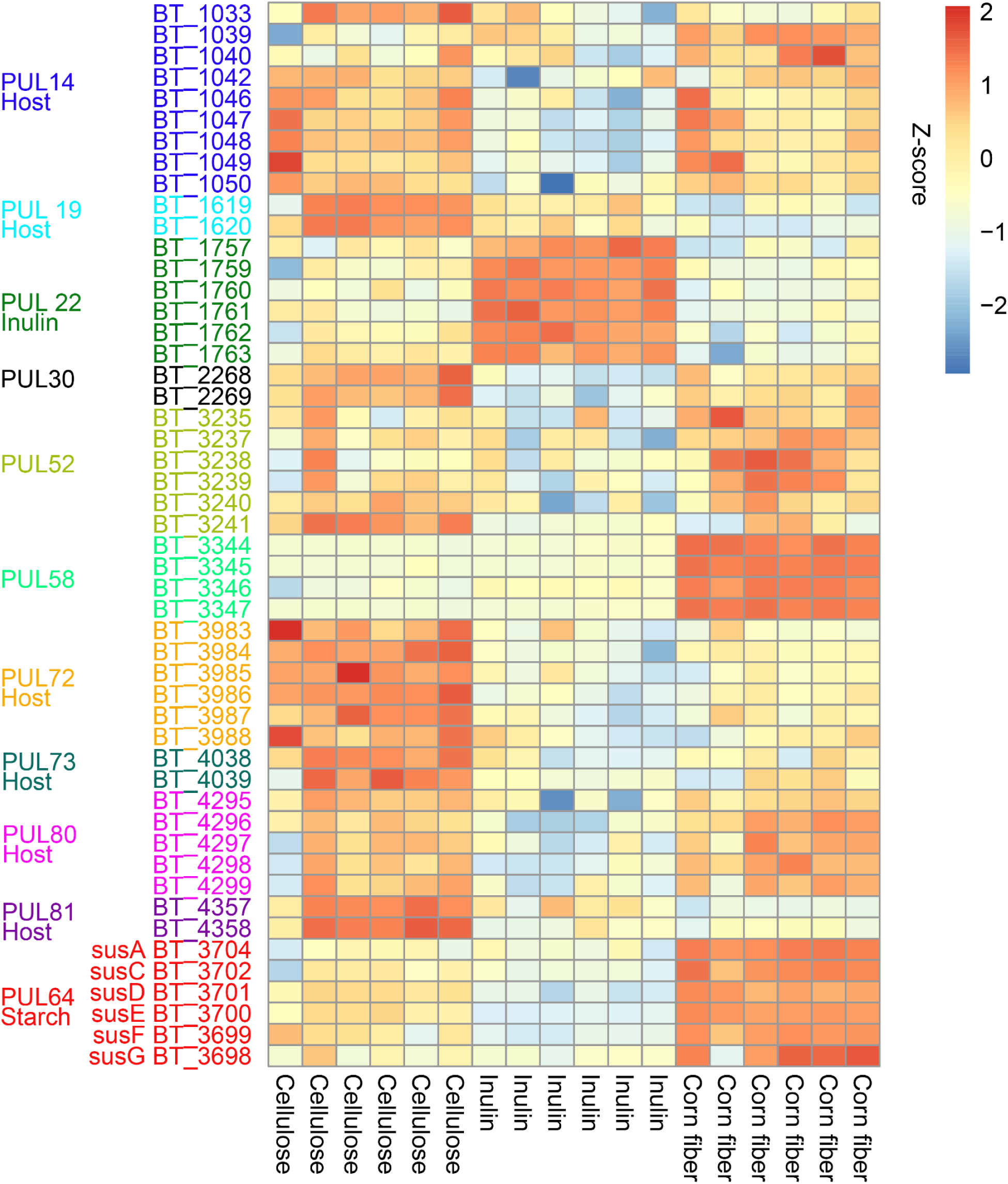
*B. thetaiotaomicron* proteins belonging to a PUL that significantly differed in abundance between the diets with different fiber sources. Heatmap ordered by diet (cellulose, inulin, corn fiber) representing the z-scored abundances of 50 proteins from PULs ordered by the PULs. If the substrate of the PUL is known, it is described. All PULs numbers are from the literature-derived numbering in PULDB

Because *B. thetaiotaomicron’s* abundance decreased under the inulin diet, its switch to inulin utilization is puzzling, given that its alternative substrates starch, host protein, and egg white dietary protein (which it used in the cellulose and corn fiber diets) were included in the inulin diet at the same amount as in the cellulose and corn fiber diets. Previous research showed that *B. thetaiotaomicron* does not grow well on inulin and that the upregulated PUL22 enables more efficient degradation of the fructan levan as compared to inulin (38). Taken together this leads us to speculate that the presence of a fructan (levan or inulin) induces expression of PUL22 and downregulates genes for the use of other carbon sources in *B. thetaiotaomicron*, whether this is beneficial for growth or not.

We also inspected the proteome of *A. muciniphila* in both Experiment 1 and the fat component of Experiment 2 because *A. muciniphila* responded with changes in I^13^C value to changes in both dietary protein and fat. Surprisingly, the abundances of only 9 proteins were significantly different in Experiment 1 (ANOVA, q <0.05) and no proteins significantly differed between fat sources in Experiment 2 (ANOVA, q <0.05), implying diet had minimal impact on *A. muciniphila’s* metabolism despite the changes in its isotopic signature in both experiments. *A. muciniphila* is known to grow almost exclusively on mucin derived from the host (12, 34) or similarly glycosylated proteins, and therefore “foraging” of host mucin could explain the changes in *A. muciniphila’s* I^13^C value in Experiment 1 (Desai et al., 2016). The change in *A. muciniphila’s* I^13^C value could, however, also be at least in part due to direct use of egg white protein as a substrate. This notion is supported by a recent study, in which the abundance of *A. muciniphila* increased when egg white was provided as the dietary protein source (Blakeley-Ruiz et al., 2024), and another study which showed that *A. muciniphila* grows on specific egg white proteins (39). In Experiment 2, the I^13^C value of host proteins was unchanged despite changes in fiber and fat source, yet A. *muciniphila’s* I^13^C value significantly decreased between corn oil and sunflower oil. This change in the I^13^C value corresponded with no change in the proteome of *A. muciniphila*.

We speculate that *A. muciniphila* consistently consumes mucin under any diet, and that different components of mucin have different isotopic signatures. Mucins are heavily glycosylated proteins, and carbon for glycosylations can come from different dietary components than carbon for the amino acids used to generate the protein backbone, which is the basis for the isotopic signature detected using protein-SIF. Specifically, sugars for glycosylations can come in part from conversion of carbohydrates and in part from gluconeogenesis with fatty acids as precursors (40). This would lead to the I^13^C value of glycosylations to change in response to changes in the isotopic signature of fiber and fat sources, while the I^13^C value of the mucin protein backbone remains unchanged as the amino acids are obtained by the host from dietary protein. Since *A. muciniphila* is predicted to be able to synthesize all the amino acids except threonine (41), it is possible that it uses carbon from mucin glycan conjugates for amino acid biosynthesis. Thus, changes in the isotopic signature of mucin glycosylations become evident in the protein I^13^C value of *A. muciniphila*. This is consistent with evidence suggesting that fat source affects glycosylations of intestinal mucins (42).

## Conclusion

In this manuscript, we used the protein-SIF approach to track the *in vivo* utilization of specific protein, fiber, and fat substrates by several species of intestinal bacteria, and quantified how utilization of these carbon sources affected accumulation of proteinaceous biomass for each bacterial species. Moreover, using the same metaproteomics data, we identified changes in the expression of particular genes that potentially underlie use of specific carbon sources and carbon source switches. Ultimately, we were able to capture 7 separate instances where a change in protein, fiber or fat source correlated with a significant change in the protein I^13^C value of a specific organism including the host, providing direct evidence for intestinal microbes using particular dietary components as carbon sources. Additionally, we found that some microbial species switch between macronutrients (protein, fiber or fat) when the source of their previously preferred macronutrient changes (Table 2).

**Table 2:**
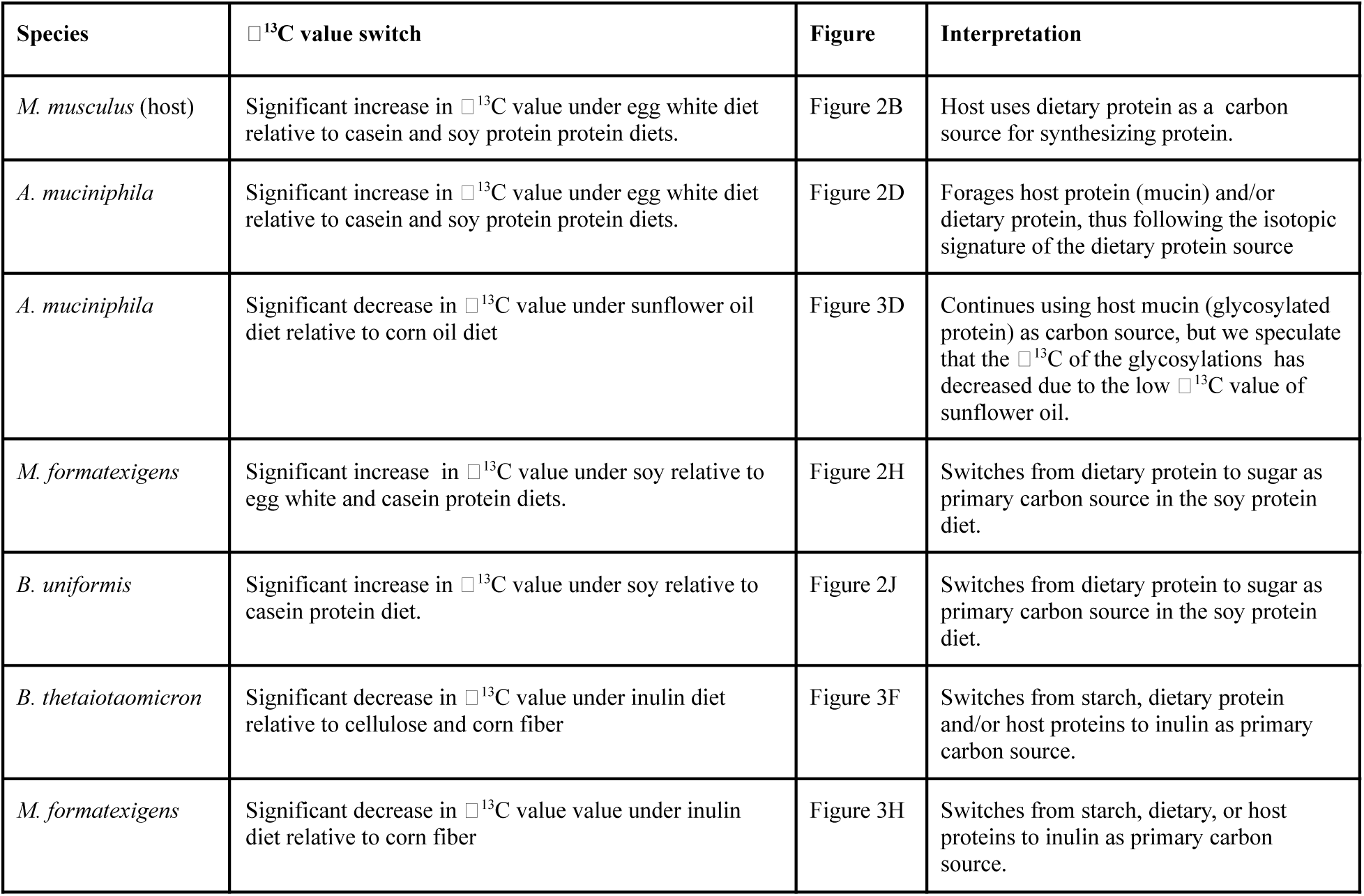
Description of significant changes in □^13^C value and their interpretation.

Our study has several limitations that could be addressed in future work. First, due to the current detection limits of the protein-SIF approach we were only able to measure I^13^C values for the more abundant members of the 13 species in the community. Currently, the speed of data acquisition of mass spectrometers is rapidly advancing and will enable calculation of I^13^C values from more community members in future experiments. We anticipate that with these advances, protein-SIF will be a powerful tool for investigating the ecology of dietary nutrient usage and niche differentiation in a variety of different host-microbe systems. Second, in this study we used gnotobiotic mice with a defined community, which has the advantage that we can use the exact protein sequences of all microbiota members to identify proteins, but it limits our understanding of how carbon sources are used in more complex, natural microbiota. We do not foresee any obstacles to applying protein-SIF to natural intestinal microbiota, particularly for the more abundant species. Third, we used fully defined diets, which are advantageous because we can precisely measure and control isotopic signatures of each diet, and also attribute changes in isotopic signature of microbial species to a specific source of the dietary component. However, fully defined diets are limited in that observed diet-microbe interactions may be driven by the “artificial”, low complexity nature of the diets. For example the unexpected switch of *B. thetaiotaomicron* to inulin utilization, which coincided with a reduction in its biomass, may not represent a realistic scenario, as the bacteria was only provided with inulin, whereas dietary fiber is usually present as a complex mixture of fiber types. In future experiments a purified food item, e.g., the prebiotic inulin, could be consumed together with regular food items with known isotopic signature (e.g. wheat bread, tofu, apple, cheese) to measure carbon source use in the context of more complex foods.. This flexibility to freely manipulate diet also highlights one of the potential advantages of protein-SIF, which is that it does not require the use of artificially labeled foods. Thus, the method could be applied to humans consuming their actual diets and without any safety concerns.

## Acknowledgments

We are sincerely grateful to Karen Flores and Dr. Susan Tonkonogy for their expertise with gnotobiotic mice and for performing all manipulations of the mice. We thank Nicholas Pudlo and Dr. Eric Martens for providing the bacterial strains. We are grateful to Deniz Durmusoglu and Dr. Nathan Crook for their assistance with growing the anaerobic strains. We thank Abigail Korenek for her assistance in preparing culture media. We thank Roxane Bowden and Dr. Christopher Osburn for the IRMS measurements of the ingredients. We thank Dr. Fernanda Salvato for providing A.M. with extensive metaproteomics training and everyone in the Kleiner Lab for discussing the experimental design and results, particularly Ali Bartlett. We made all LC-MS/MS measurements in the Molecular Education, Technology, and Research Innovation Center (METRIC) at NC State University.

## Declarations

### Ethics Approval

The protocols for husbandry and experimentation of all mice used in this study were approved by the Institutional Animal Care and Use Committee at North Carolina State University (Institution reference: D16-00214).

### Availability of Data

The mass spectrometry metaproteomics data and protein sequence database were deposited to the ProteomeXchange Consortium via the PRIDE (43) partner repository. PXD046928.

## References

1. B. K. Perler, E. S. Friedman, G. D. Wu, The Role of the Gut Microbiota in the Relationship Between Diet and Human Health. Annu. Rev. Physiol. 85, 449–468 (2023).

2. S. Kim, et al., Mucin degrader Akkermansia muciniphila accelerates intestinal stem cell-mediated epithelial development. Gut Microbes 13, 1892441 (2021).

3. A. Shimotoyodome, S. Meguro, T. Hase, I. Tokimitsu, T. Sakata, Short chain fatty acids but not lactate or succinate stimulate mucus release in the rat colon. Comp. Biochem. Physiol. A. Mol. Integr. Physiol. 125, 525–531 (2000).

4. M. Sun, et al., Microbiota-derived short-chain fatty acids promote Th1 cell IL-10 production to maintain intestinal homeostasis. Nat. Commun. 9, 3555 (2018).

5. A. Bartlett, M. Kleiner, Dietary protein and the intestinal microbiota: An understudied relationship. iScience 25, 105313 (2022).

6. K. Oliphant, E. Allen-Vercoe, Macronutrient metabolism by the human gut microbiome: major fermentation by-products and their impact on host health. Microbiome 7, 91 (2019).

7. L. A. David, et al., Diet rapidly and reproducibly alters the human gut microbiome. Nature 505, 559–563 (2014).

8. J. J. Faith, N. P. McNulty, F. E. Rey, J. I. Gordon, Predicting a Human Gut Microbiota’s Response to Diet in Gnotobiotic Mice. Science 333, 101–104 (2011).

9. Y. Zhu, et al., Meat, dairy and plant proteins alter bacterial composition of rat gut bacteria. Sci. Rep. 5, 15220 (2015).

10. P. Li, et al., Systematic evaluation of antimicrobial food preservatives on glucose metabolism and gut microbiota in healthy mice. Npj Sci. Food 6, 42 (2022).

11. E. J. Culp, A. L. Goodman, Cross-feeding in the gut microbiome: Ecology and mechanisms. Cell Host Microbe 31, 485–499 (2023).

12. M. S. Desai, et al., A Dietary Fiber-Deprived Gut Microbiota Degrades the Colonic Mucus Barrier and Enhances Pathogen Susceptibility. Cell 167, 1339–1353.e21 (2016).

13. J. L. Sonnenburg, et al., Glycan Foraging in Vivo by an Intestine-Adapted Bacterial Symbiont. Science 307, 1955–1959 (2005).

14. E. C. Martens, H. C. Chiang, J. I. Gordon, Mucosal glycan foraging enhances fitness and transmission of a saccharolytic human gut bacterial symbiont. Cell Host Microbe 4, 447–457 (2008).

15. H. Liu, et al., Functional genetics of human gut commensal Bacteroides thetaiotaomicron reveals metabolic requirements for growth across environments. Cell Rep. 34 (2021).

16. N. Crook, et al., Adaptive Strategies of the Candidate Probiotic E. coli Nissle in the Mammalian Gut. Cell Host Microbe 25, 499–512.e8 (2019).

17. X. Zeng, et al., Gut bacterial nutrient preferences quantified in vivo. Cell 185, 3441–3456.e19 (2022).

18. M. J. DeNiro, S. Epstein, Influence of diet on the distribution of carbon isotopes in animals. Geochim. Cosmochim. Acta 42, 495–506 (1978).

19. M. Kleiner, et al., Metaproteomics method to determine carbon sources and assimilation pathways of species in microbial communities. Proc. Natl. Acad. Sci. 115, E5576–E5584 (2018).

20. M. Kleiner, et al., Ultra-sensitive isotope probing to quantify activity and substrate assimilation in microbiomes. Microbiome 11, 24 (2023).

21. V. J. Orphan, C. H. House, K.-U. Hinrichs, K. D. McKeegan, E. F. DeLong, Methane-Consuming Archaea Revealed by Directly Coupled Isotopic and Phylogenetic Analysis. Science 293, 484–487 (2001).

22. A. Pearson, “Pathways of Carbon Assimilation and Their Impact on Organic Matter Values δ13C” in Handbook of Hydrocarbon and Lipid Microbiology, K. N. Timmis, Ed. (Springer Berlin Heidelberg, 2010), pp. 143–156.

23. D. Berry, et al., Host-compound foraging by intestinal microbiota revealed by single-cell stable isotope probing. Proc. Natl. Acad. Sci. 110, 4720 (2013).

24. A. T. Reese, et al., Microbial nitrogen limitation in the mammalian large intestine. Nat. Microbiol. 3, 1441–1450 (2018).

25. M. L. Patnode, et al., Interspecies Competition Impacts Targeted Manipulation of Human Gut Bacteria by Fiber-Derived Glycans. Cell 179, 59–73.e13 (2019).

26. J. A. Blakeley-Ruiz, et al., Dietary protein source strongly alters gut microbiota composition and function. bioRxiv 2024.04.04.588169 (2024). 10.1101/2024.04.04.588169.

27. A. Mordant, M. Kleiner, Evaluation of Sample Preservation and Storage Methods for Metaproteomics Analysis of Intestinal Microbiomes. Microbiol. Spectr. 9, e0187721 (2021).

28. P. G. Reeves, F. H. Nielsen, G. C. Fahey, AIN-93 Purified Diets for Laboratory Rodents: Final Report of the American Institute of Nutrition Ad Hoc Writing Committee on the Reformulation of the AIN-76A Rodent Diet. J. Nutr. 123, 1939–1951 (1993).

29. J. R. Wiśniewski, A. Zougman, N. Nagaraj, M. Mann, Universal sample preparation method for proteome analysis. Nat. Methods 6, 359–362 (2009).

30. J. A. Blakeley-Ruiz, M. Kleiner, Considerations for constructing a protein sequence database for metaproteomics. Comput. Struct. Biotechnol. J. 20, 937–952 (2022).

31. The UniProt Consortium, UniProt: the Universal Protein Knowledgebase in 2023. Nucleic Acids Res. 51, D523–D531 (2023).

32. M. Kleiner, et al., Assessing species biomass contributions in microbial communities via metaproteomics. Nat. Commun. 8, 1558 (2017).

33. S. Tyanova, et al., The Perseus computational platform for comprehensive analysis of (prote)omics data. Nat. Methods 13, 731–740 (2016).

34. M. Derrien, E. E. Vaughan, C. M. Plugge, W. M. de Vos, Akkermansia muciniphila gen. nov., sp. nov., a human intestinal mucin-degrading bacterium. Int. J. Syst. Evol. Microbiol. 54, 1469–1476 (2004).

35. L. Reitzer, “Amino Acid Synthesis” in Encyclopedia of Microbiology (Third Edition), M. Schaechter, Ed. (Academic Press, 2009), pp. 1–17.

36. J. M. Grondin, K. Tamura, G. Déjean, D. W. Wade, H. Brumer, Polysaccharide Utilization Loci: Fueling Microbial Communities. J. Bacteriol. 199, 10.1128/jb.00860-16 (2017).

37. N. Terrapon, et al., PULDB: the expanded database of Polysaccharide Utilization Loci. Nucleic Acids Res. 46, D677–D683 (2018).

38. E. D. Sonnenburg, et al., Specificity of Polysaccharide Use in Intestinal Bacteroides Species Determines Diet-Induced Microbiota Alterations. Cell 141, 1241–1252 (2010).

39. H. Takada, T. Katoh, T. Katayama, Sialylated O -Glycans from Hen Egg White Ovomucin are Decomposed by Mucin-degrading Gut Microbes. J. Appl. Glycosci. 67, 31–39 (2020).

40. M. H. Green, Are Fatty Acids Gluconeogenic Precursors? J. Nutr. 150, 2235–2238 (2020).

41. Ottman Noora, et al., Genome-Scale Model and Omics Analysis of Metabolic Capacities of Akkermansia muciniphila Reveal a Preferential Mucin-Degrading Lifestyle. Appl. Environ. Microbiol. 83, e01014–17 (2017).

42. M. Mastrodonato, G. Calamita, D. Mentino, G. Scillitani, High-fat Diet Alters the Glycosylation Patterns of Duodenal Mucins in a Murine Model. J. Histochem. Cytochem. 68, 279–294 (2020).

43. Y. Perez-Riverol, et al., The PRIDE database resources in 2022: a hub for mass spectrometry-based proteomics evidences. Nucleic Acids Res. 50, D543–D552 (2022).

