## Supplementary Materials for "Stable isotope fingerprinting can directly link intestinal microorganisms with their carbon source and captures diet-induced substrate switching *in vivo*"

**Supplementary Table 1:** Diet formulations.

| Teklad ID | TD.200545 | TD.200546 | TD.200547 | TD.200548 | TD.200549 | TD.200550 | TD.200551 |
|---|---|---|---|---|---|---|---|
| Name | Egg white protein, corn oil, soluble corn fiber diet | Cellulose fiber diet | Inulin fiber diet | Sunflower oil diet | Casein protein diet | Soy protein diet | Soybean oil diet |
| Protein | Egg white 200g/Kg | Egg white 200g/Kg | Egg white 200g/Kg | Egg white 200g/Kg | Casein 200g/Kg | Soy 200g/Kg | Egg white 200g/Kg |
| Corn starch | 351.371 g/Kg | 351.371 g/Kg | 351.371 g/Kg | 351.371 g/Kg | 351.371 g/Kg | 351.371 g/Kg | 351.371 g/Kg |
| Maltodextrin | 132 g/Kg | 132 g/Kg | 132 g/Kg | 132 g/Kg | 132 g/Kg | 132 g/Kg | 132 g/Kg |
| Sucrose | 100 g/Kg | 100 g/Kg | 100 g/Kg | 100 g/Kg | 100 g/Kg | 100 g/Kg | 100 g/Kg |
| oil | Corn oil 120 g/Kg | Corn oil 120 g/Kg | Corn oil 120 g/Kg | Sunflower oil 120 g/Kg | Corn oil 120 g/Kg | Corn oil 120 g/Kg | Soybean oil 120 g/Kg |
| fiber | Soluble corn fiber 50 g/Kg | Cellulose 50 g/Kg | Inulin 50 g/Kg | Soluble corn fiber 50 g/Kg | Soluble corn fiber 50 g/Kg | Soluble corn fiber 50 g/Kg | Soluble corn fiber 50 g/Kg |
| Mineral mix | 13.39 g/Kg | 13.39 g/Kg | 13.39 g/Kg | 13.39 g/Kg | 13.39 g/Kg | 13.39 g/Kg | 13.39 g/Kg |
| Calcium phosphate | 12.02 g/Kg | 12.02 g/Kg | 12.02 g/Kg | 12.02 g/Kg | 12.02 g/Kg | 12.02 g/Kg | 12.02 g/Kg |
| Calcium Carbonate | 3.43 g/Kg | 3.43 g/Kg | 3.43 g/Kg | 3.43 g/Kg | 3.43 g/Kg | 3.43 g/Kg | 3.43 g/Kg |
| Vitamin mix | 15 g/Kg | 15 g/Kg | 15 g/Kg | 15 g/Kg | 15 g/Kg | 15 g/Kg | 15 g/Kg |
| Choline | 2.75 g/Kg | 2.75 g/Kg | 2.75 g/Kg | 2.75 g/Kg | 2.75 g/Kg | 2.75 g/Kg | 2.75 g/Kg |
| Biotin | 0.003 g/Kg | 0.003 g/Kg | 0.003 g/Kg | 0.003 g/Kg | 0.003 g/Kg | 0.003 g/Kg | 0.003 g/Kg |
| Thiamin | 0.01 g/Kg | 0.01 g/Kg | 0.01 g/Kg | 0.01 g/Kg | 0.01 g/Kg | 0.01 g/Kg | 0.01 g/Kg |
| Vitamin K | 0.002 g/Kg | 0.002 g/Kg | 0.002 g/Kg | 0.002 g/Kg | 0.002 g/Kg | 0.002 g/Kg | 0.002 g/Kg |
| TBHQ | 0.024 g/Kg | 0.024 g/Kg | 0.024 g/Kg | 0.024 g/Kg | 0.024 g/Kg | 0.024 g/Kg | 0.024 g/Kg |

**Supplementary Table 2. EA-IRMS results of dietary components used in this study**. Unless indicated, all ingredient samples were obtained from Envigo Teklad. Duplicate dietary component rows indicate duplicate measurements.

| Dietary component | mg C | mg N | $\delta^{13}C$ | $\delta^{15}N$ | C:N | Avg. $\delta^{13}C$ (n=2) | Diff. reps (n = 2) | Date tested |
|---|---|---|---|---|---|---|---|---|
| cornstarch | 0.073 | | -10.84 | | | **-10.75** | 0.13 | Jan 10 2019 |
| cornstarch | 0.122 | | -10.66 | | | | | Jan 10 2019 |
| sucrose | 0.061 | | -12.18 | | | **-12.32** | 0.19 | Jan 10 2019 |
| sucrose | 0.204 | | -12.46 | | | | | Jan 10 2019 |
| cellulose | 0.193 | | -26.41 | | | **-26.55** | 0.20 | Jan 10 2019 |
| cellulose | 0.144 | | -26.69 | | | | | Jan 10 2019 |
| inulin | 0.125 | | -26.92 | | | **-26.88** | 0.06 | Jan 10 2019 |
| inulin | 0.092 | | -26.83 | | | | | Jan 10 2019 |
| maltodextrin maltrin | 0.238 | | -10.40 | | | **-10.40** | 0.01 | Jan 10 2019 |
| maltodextrin maltrin | 0.172 | | -10.39 | | | | | Jan 10 2019 |
| maltodextrin lodex | 0.166 | | -10.58 | | | **-10.65** | 0.09 | Jan 10 2019 |
| maltodextrin lodex | 0.101 | | -10.71 | | | | | Jan 10 2019 |
| casein | 0.091 | 0.02829 | -26.46 | 6.02 | 3.7 | **-26.56** | 0.15 | Jan 10 2019 |
| casein | 0.095 | 0.02832 | -26.67 | 5.19 | 3.9 | | | Jan 10 2019 |
| soy protein | 0.112 | 0.03342 | -26.36 | -0.22 | 3.9 | **-26.33** | 0.05 | Jan 10 2019 |
| soy protein | 0.070 | 0.02152 | -26.29 | -0.06 | 3.8 | | | Jan 10 2019 |
| egg white solids | 0.147 | 0.04316 | -17.03 | 4.31 | 4.0 | **-17.19** | 0.22 | Jan 10 2019 |
| egg white solids | 0.113 | 0.03324 | -17.35 | 4.05 | 4.0 | | | Jan 10 2019 |
| soybean oil | 0.137 | | -32.06 | | | **-32.05** | 0.01 | Jan 10 2019 |
| soybean oil | 0.315 | | -32.05 | | | | | Jan 10 2019 |
| corn oil | 0.142 | | -17.16 | | | **-16.92** | 0.35 | Jan 10 2019 |
| corn oil | 0.240 | | -16.67 | | | | | Jan 10 2019 |
| Corn fiber Amazon B00NAD0IVU | 0.091 | 0.0005 | -11.11 | | | **-11.17** | 0.07 | Apr 2019 |
| Corn fiber Amazon B00NAD0IVU | 0.180 | 0.0005 | -11.15 | | | | | Apr 2019 |
| Corn fiber Amazon B00NAD0IVU | 0.255 | 0.0005 | -11.24 | | | | | Apr 2019 |
| Sunflower oil Amazon B0792GCNWV | 0.148 | | -30.95 | | | **-31.24** | 0.42 | Apr 2019 |
| Sunflower oil Amazon B0792GCNWV | 0.240 | | -31.54 | | | | | Apr 2019 |

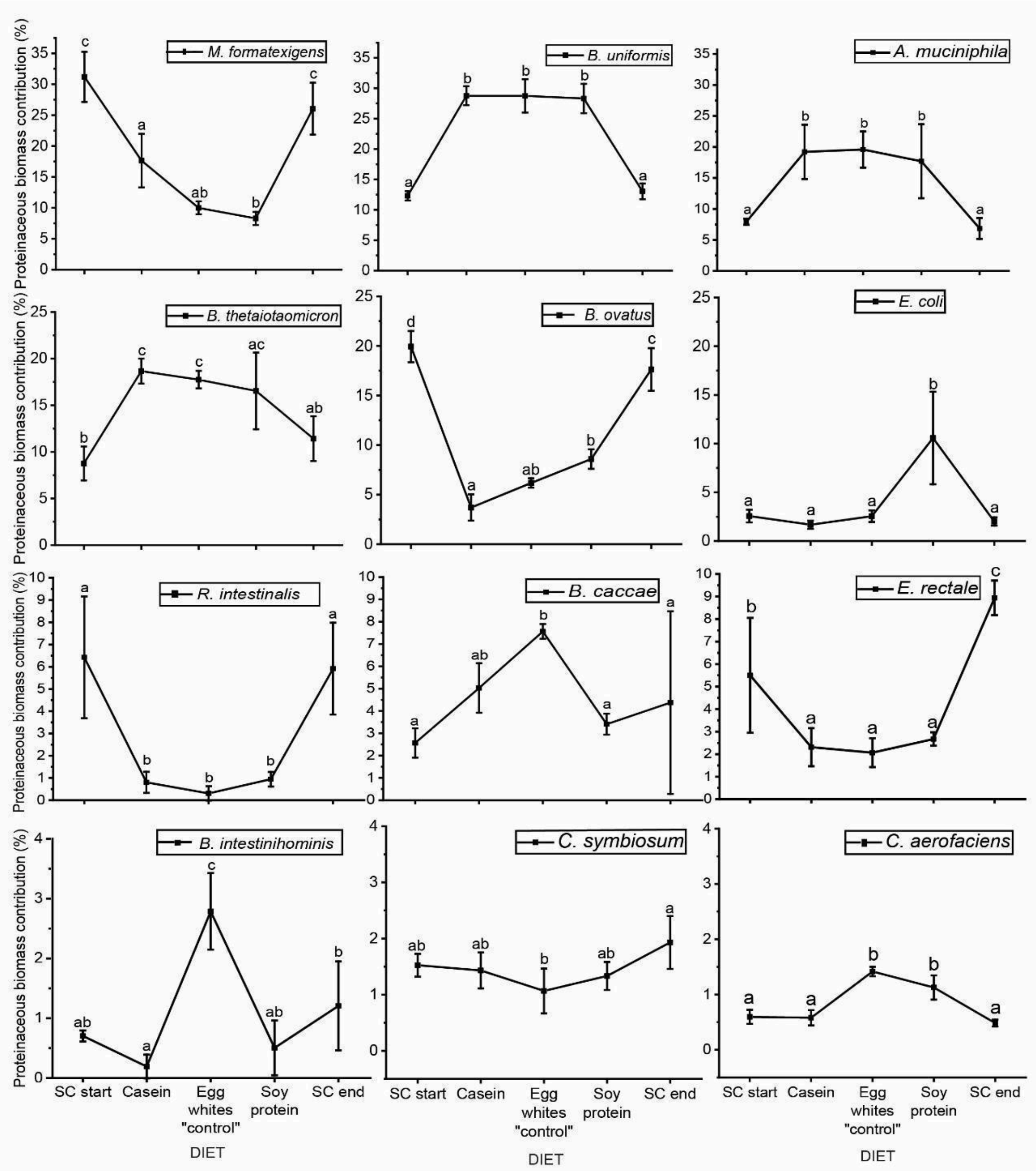


**Supplementary Figure 1. Relative proteinaceous biomass contribution of *M. formatexigens, B. uniformis, A. muciniphila, B. thetaiotaomicron, B. ovatus, E. coli, R. intestinalis, B. caccae, E. rectale, B. intestinihominis, C. symbiosum, and C. aerofaciens* in mice fed the protein diets (Experiment 1).** Each point represents the relative biomass contribution of the organism after the mice were fed the diet indicated on the x-axis for seven days. Relative abundances were averaged and error bars indicate standard deviation (n = 5). Diets are ordered on the x-axis in chronological order fed to the mice. SC = standard chow diet. Casein = diet with Casein as the protein source; Egg whites ("control") = diet with egg whites as the protein source; Soy protein = diet with soy protein as the protein source. Please note that the y-axis scale differs per row. Significant differences are indicated by different letters (a, b, c, d; based on one-way ANOVA and Tukey's HSD post hoc test, $p < .05$).

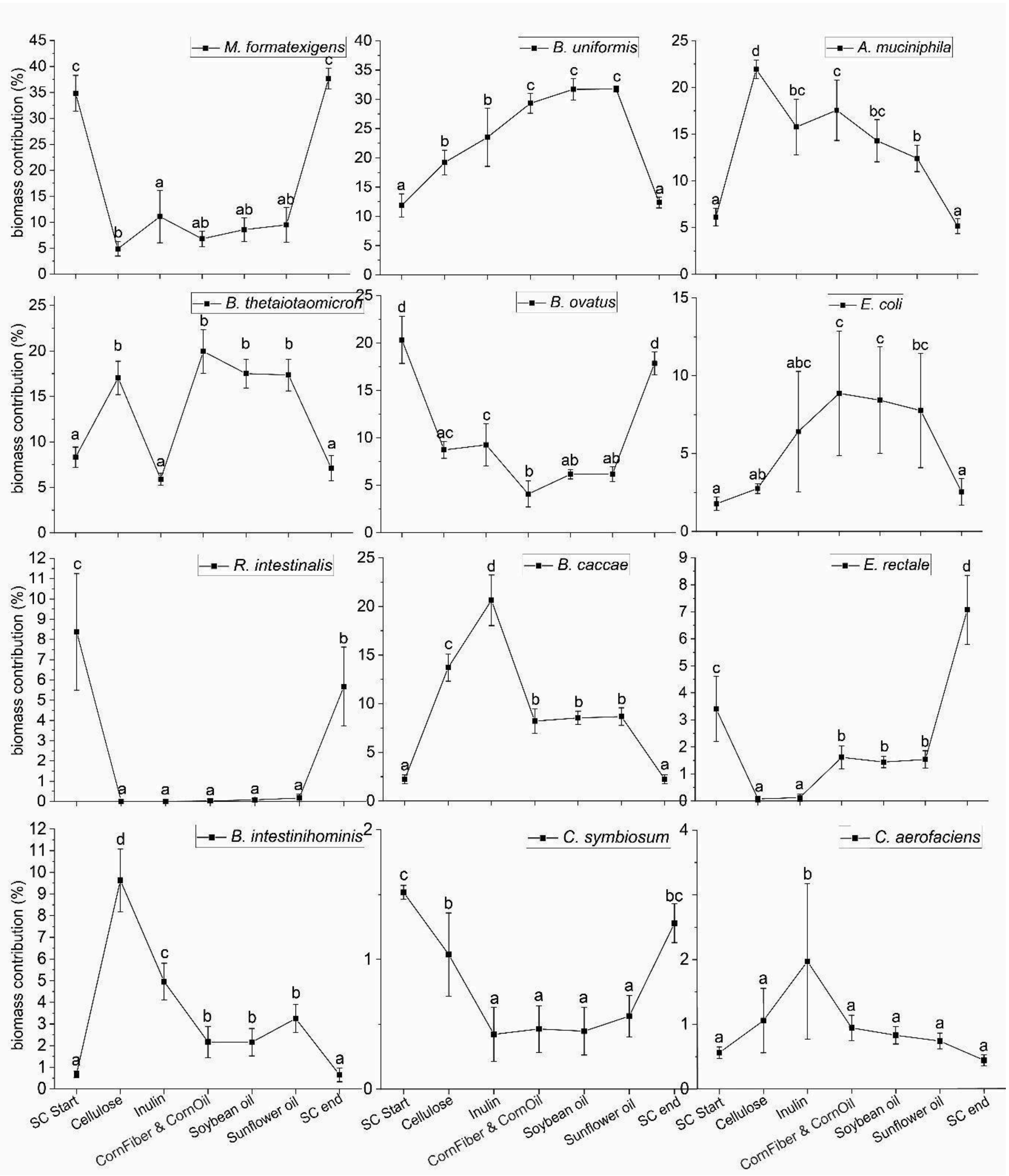


**Supplementary Figure 2. Relative proteinaceous biomass contribution of *M. formatexigens, B. uniformis, A. muciniphila, B. thetaiotaomicron, B. ovatus, E. coli, R. intestinalis, B. caccae, E. rectale, B. intestinihominis, C. symbiosum*, and *C. aerofaciens* in mice fed the fiber and fat diets (Experiment 2).** Each point represents the relative biomass contribution of the organism after the mice were fed the diet indicated on the x-axis for seven days. Relative abundances were averaged and error bars indicate standard deviation (n = 6). Diets are ordered on the x-axis in chronological order fed to the mice. SC = standard chow diet. Cellulose = diet with cellulose as the fiber source; Corn fiber & Corn oil ("control") = diet with corn fiber as the fiber source and corn oil as the fat source; Soybean oil = diet with soybean oil as the fat source. Sunflower oil = diet with sunflower oil as the fat source. Please note that the y-axis scale differs per row. Significant differences are indicated by different letters (a, b, c, d; based on one-way ANOVA and Tukey's HSD post hoc test, $p < .05$).
